# Diffantom: whole-brain diffusion MRI phantoms derived from real datasets of the Human Connectome Project

**DOI:** 10.1101/026898

**Authors:** Oscar Esteban, Emmanuel Caruyer, Alessandro Daducci, Meritxell Bach-Cuadra, Maria-J. Ledesma-Carbayo, Andres Santos

**Author notes:** Corresponding Author: Oscar Esteban Biomedical Image Technologies (BIT), ETSI Telecomunicación, Av. Complutense 30, C203, E28040 Madrid, Spain.

## Abstract

Diffantom is a whole-brain digital phantom generated from a dataset from the Human Connectome Project. Diffantom is presented here to be openly and freely distributed along with the diffantomizer workflow to generate new diffantoms. We encourage the neuroimage community to contribute with their own diffantoms and share them openly.

## INTRODUCTION

Fiber tracking on diffusion MRI (dMRI) data has become an important tool for the *in-vivo* investigation of the structural configuration of fiber bundles at the macroscale. Tractography is fundamental to gain information about white matter (WM) morphology in many clinical applications like neurosurgical planning (Golby et al., 2011), post-surgery evaluations (Toda et al., 2014), and the study of neurological diseases as in (Chua et al., 2008) addressing multiple sclerosis and Alzheimer’s disease. The analysis of structural brain networks using graph theory is also applied on tractography, for instance in the definition of the unique subject-wise patterns of connectivity (Sporns et al., 2005), in the assessment of neurological diseases (Griffa et al., 2013), and in the study of the link between structural and functional connectivity (Messé et al., 2015). However, the development of the field is limited by the lack of a gold standard to test and compare the wide range of methodologies available for processing and analyzing dMRI.

Large efforts have been devoted to the development of physical phantoms (Lin et al., 2001; Campbell et al., 2005; Perrin et al., 2005; Fieremans et al., 2008; Tournier et al., 2008). Côté et al. (2013) conducted a thorough review of tractography methodologies using the so-called *FiberCup* phantom (Poupon et al., 2008; Fillard et al., 2011). These phantoms are appropriate to evaluate the angular resolution in fiber crossings and accuracy of direction-independent scalar parameters in very simplistic geometries. Digital simulations are increasingly popular because the complexity of whole-brain tractography can not be accounted for with current materials and proposed methodologies to build physical phantoms. Early digital phantoms started with simulation of simple geometries (Basser et al., 2000; Gössl et al., 2002; Tournier et al., 2002; Leemans et al., 2005) to evaluate the angular resolution as well. These tools generally implemented the multi-tensor model (Alexander et al., 2001; Tuch et al., 2002) to simulate fiber crossing, fanning, kissing, etc. Close et al. (2009) presented the *Numerical Fibre Generator*, a software to simulate spherical shapes filled with digital fiber tracts. Caruyer et al. (2014) proposed *Phantomas* to simulate any kind of analytic geometry inside a sphere. *Phantomas* models diffusion by a restricted and a hindered compartment, similar to (Assaf and Basser, 2005). Wilkins et al. (2014) proposed a whole-brain simulated phantom derived from voxel-wise orientation of fibers averaged from real dMRI scans and the multi-tensor model with a compartment of isotropic diffusion. Neher et al. (2014) proposed *FiberFox*, a visualization software to develop complex geometries and their analytical description. Once the geometries are obtained, the software generates the corresponding dMRI signal with a methodology very close to that implemented in *Phantomas*. An interesting outcome of *FiberFox* is the phantom dataset^1^ created for the Tractography Challenge held in ISMRM 2015. This dataset was derived from the tractography extracted in one Human Connectome Project (HCP, Van Essen et al. (2012)) dataset. In the tractogram, 25 fiber bundles of interest were manually segmented by experts. Using *FiberFox*, the segmentation of each bundle was mapped to an analytical description, and finally simulated the signal.

In this data report we present *Diffantom*, an *in-silico* dataset to assess tractography and connectivity pipelines using dMRI real data as source microstructural information. *Diffantom* is inspired by the work of Wilkins et al. (2014), with two principal novelties. First, since we use a dataset from the HCP as input, data are already corrected for the most relevant distortions. The second improvement is a more advanced signal model to generate the phantom using the hindered and restricted diffusion model of *Phantomas* (Caruyer et al., 2014). As a result, we provide a whole-brain digital phantom of dMRI data with structural information derived from an HCP dataset. We also openly release the *diffantomizer* workflow, the software package necessary to generate custom *diffantoms*. *Diffantom* is originally designed for the investigation of susceptibility-derived distortions, a typical artifact that produces geometrical warping in certain regions of dMRI datasets. In (Esteban et al., 2014) we addressed this phenomenon and concluded that the connectivity matrix of *Phantomas* was not dense enough to evaluate the integration of correction methods in pipelines for the connectome extraction.

## DATA DESCRIPTION

### Microstructural model

The simulation process relies on a microstructural model derived from real data. On one hand, the *diffantomizer* workflow requires up to five fraction maps {*T_j_ | j ∈* { 1, *…*, 5}} of free-and hindered-diffusion (see Figure 1, panel A). These compartments will be derived from the macroscopic structure of tissues within the brain, specified in the following order^2^: cortical gray matter (cGM), deep gray matter (dGM), WM, cerebrospinal fluid (CSF), and abnormal tissue^3^. On the other hand, the restricted-diffusion compartments are specified by up to three volume fractions {*F_i_ | i ∈* {1, 2, 3}} of three single fiber populations per voxel along with their corresponding direction maps {**V**_*i*_ *| i ∈* {1, 2, 3}}.

**Figure 1.**
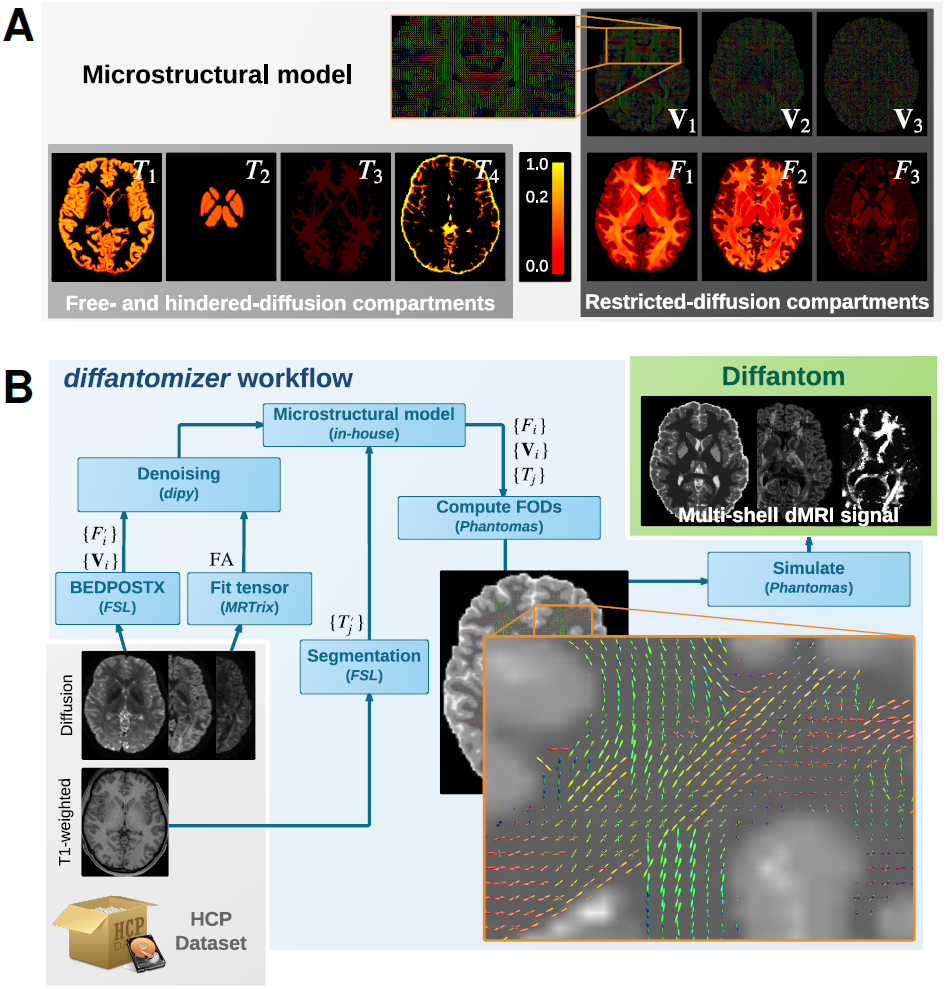
**A. Microstructural model of *Diffantom***. The phantom is simulated from an underlying microstructural model specified with the following volume-fraction maps: three hindered-diffusion compartments {*T*_1_, *T*_2_, *T*_3_}, one free-diffusion compartment *T*_4_ corresponding to the cerebrospinal fluid (CSF), three restricted-diffusion compartments {*F*_*i*_}, and three vectorial maps associated with the local fiber directions {**V**_*i*_}. Please note the piece-wise linear function of the color scale to enable visibility of small volume fractions. **B. The *diffantomizer* workflow, a workflow to generate *diffantoms***. The pipeline to generate phantoms from any Human Connectome Project (HCP, Van Essen et al. (2012)) dataset is presented in the lower panel. Once the microstructural model shown in the upper panel has been prepared as described in Data description, the local orientations are computed and fed into *Phantomas* to finally simulate the signal.

The process to obtain the microstructural model from one dataset of the HCP can be described as follows (see also Figure 1, panel B): 1) The fiber orientation maps {**V**_*i*_} and their corresponding estimations of volume fraction 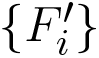 are obtained using the ball-and-stick model for multi-shell data of BEDPOSTX (Bayesian Estimation of Diffusion Parameters Obtained using Sampling Techniques modelling crossing –X– fibres, Jbabdi et al. (2012)) on the dMRI data. The HCP recommends BEDPOSTX to reconstruct their data (Glasser et al., 2013). A further advantage is that BEDPOSTX exploits the multi-shell acquisitions of the HCP while operating at whole-brain level. 2) A fractional anisotropy (FA) map is obtained after fitting a tensor model with *MRTrix*. As we shall see in the Appendix, the FA is used to infer *F*_1_ (the fraction map of the most prevalent fiber), avoiding the extremely noisy estimation of 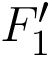 performed by BEDPOSTX in the previous step. 3) The original fiber fractions 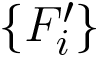 and the FA map are denoised with a nonlocal means filter included in *dipy* (Garyfallidis et al., 2014). This step produces an important smoothing of the maps, while preserving the edges. Smoothing is also beneficial in simplifying the voxel-wise diffusion model. 4) The macrostructural fractions 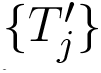 are extracted from the T1-weighted image of the dataset, using standard *FSL* segmentation tools (Jenkinson et al., 2012). 5) The images obtained previously (FA map, {**V**_*i*_}, 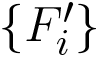, and 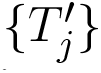) are combined as described in the Appendix to generate the final microstructural model ({**V**_*i*_}, {*F_i_*}, and {*T_j_*}), presented in Figure 1-A.

### Diffusion signal generation

Once a microstructural model of the subject has been synthesized, the fiber orientation maps {**V**_*i*_} are weighted by the fiber-fraction maps {*F*_*i*_} and projected onto a continuous representation of the fiber orientation distributions (FODs). A close-up showing how the FODs map looks is presented in Figure 1B. The single fiber response is a Gaussian diffusion tensor with axial symmetry and eigenvalues *λ*_1_ = 2.2 · 10^-3^ mm^2^s^-1^ and *λ*_2,3_ = 0.2 · 10^-3^ mm^2^s^−1^. The resulting FODs map is then combined with the free-and hindered-diffusion compartments corresponding to {*T_j_*}. The free-diffusion compartment corresponds to the CSF fraction map *T*_4_ and is modeled with isotropic diffusivity *D*_*CSF*_ of 3.0 · 10^-3^ mm^2^s^-1^. The hindered-diffusion compartments correspond to {*T*_1_, *T*_2_, *T*_3}_ and are also modeled with isotropic diffusivity *D*_*W* *M*_ = 2.0 · 10^-4^, *D*_*cGM*_ = 7.0 · 10^-4^ and *D*_*dGM*_ = 9.0 · 10^-4^, respectively [mm^2^s^-1^]. All these values for diffusivity (and the corresponding to the single-fiber response) can be modified by the user with custom settings. The restricted-and hindered-compartments are then fed into *Phantomas* (Caruyer et al., 2014) and the final dMRI signal is obtained. By default, diffusion data are generated using a scheme of 100 directions distributed in one shell with uniform coverage (Caruyer et al., 2013). Custom one-or multi-shell schemes can be generated supplying the tables of corresponding vectors and *b*-values. Rician noise is also included in *Phantomas*, and the signal-to-noise ratio (SNR) can be set by the user. The default value for SNR is preset to 30.0.

### Implementation and reproducibility

We also provide the *diffantomizer* workflow, the software package used to generate *diffantoms*, so that users can regenerate similar datasets with different parameters. This workflow, presented in Figure 1, is implemented using *nipype* (Gorgolewski et al., 2011) to ensure reproducibility and usability.

### Interpretation and recommended uses

To illustrate the features of *Diffantom*, the example dataset underwent a simplified connectivity pipeline including constrained spherical deconvolution (CSD) and probabilistic tractography from *MRTrix* (Tournier et al., 2012). CSD was reconstructed using 8^th^-order spherical harmonics, and tractography with 1.6 · 10^6^ seed points evenly distributed across a dilated mask of the WM tissue. Figure 2, panels A1 and A3, show the result of the tractography obtained with such pipeline for the original *Diffantom* and a distorted version. Finally, we applied *tract querier* (Wassermann et al., 2013) to segment some fiber bundles such as the corticospinal tract (CST) and the forceps minor (see Figure 2, panels A2, A4). Particularly, due to its location nearby the orbitofrontal lobe, the forceps minor is generally affected by susceptibility distortions.

**Figure 2.**
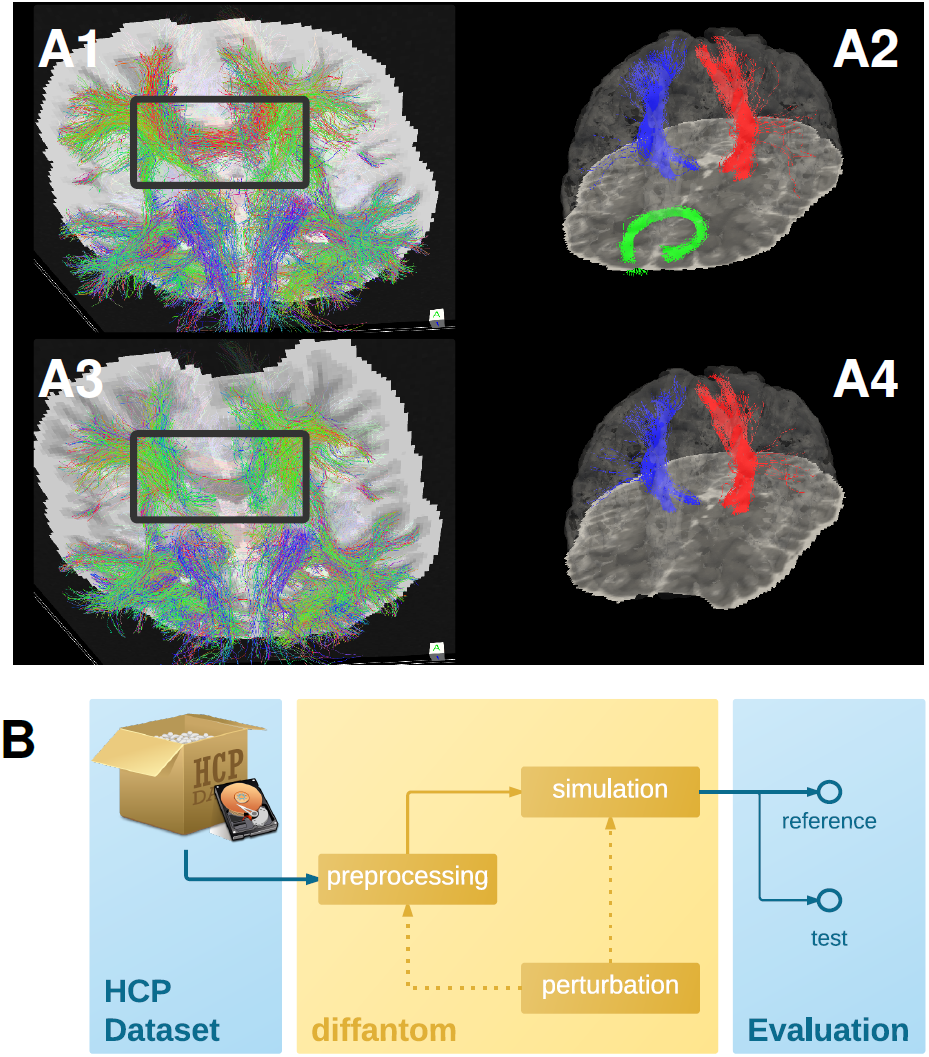
**A. Example dataset.** A1 and A3 show the tractogram of fibers crossing slice 56 of *Diffantom* as extracted with *MRTrix*, represented over the corresponding slice of the *b0* volume for the original (A1) and the distorted (A3) phantoms, with a gray frame highlighting the absence of important tracks. Panels A2 and A4 show the segmentation of the right corticospinal tract (CST) represented with blue streamlines, the left CST (red streamlines), and the forceps minor (green streamlines) using *tract querier*. A2 and A4 include the slice 56 of the *b0* and the pial surface is represented with transparency. In the distorted *Diffantom* (A4) the forceps minor was not detected. **B. Recommended use of *Diffantom***. The phantom is designed to be used as ground-truth information in evaluation frameworks, to implement unit test of algorithms, to check integration of processing units within pipelines or to validate complete workflows. For instance, in order to evaluate artifacts, a perturbation can be induced in the microstructural model or after simulation to provide <u>reference and test datasets.</u>

We recommend *Diffantom* as ground-truth in verification and validation frameworks (Figure 2, panel B) for testing pipelines. *Diffantom* is applicable in the unit testing of algorithms, the integration testing of modules in workflows, and the overall system testing. Some potential applications follow:

- Investigating the impact of different diffusion sampling schemes on the local microstructure model of choice and on the subsequent global tractography outcome. Since the gradient scheme can be set by the user, *Diffantom* can be seen as a mean to translate the so-called *b-matrix* of the source dataset to any target scheme.
- Assessment of sensitivity and robustness to imaging artifacts (noise, partial volume effect and CSF contamination, susceptibility-derived warping, Eddy-currents-derived distortions, etc.) at unit, integration and systems testing levels.
- Using *Diffantom* as in panel B of Figure 2, it is possible to apply binary classification measures to evaluate the resulting connectivity matrix. Considering the connectivity matrix of the *reference Diffantom* and the resulting matrix of the *test Diffantom*, the receiver operating characteristic (ROC) of the pipeline can be characterized.
- Simulation of pathological brains by altering the microstructural model accordingly (e.g. as tumors were simulated in Kaus et al. (2000)).

In order to exemplify one of these intended uses, we also release a *Diffantom* including the susceptibility derived distortion in simulation.

## DISCUSSION AND CONCLUSION

### Discussion

Whole-brain, realistic dMRI phantoms are necessary in the developing field of structural connectomics. *Diffantom* is a derivative of (Wilkins et al., 2014) in terms of methodology for simulation with two major advances. First, the correctness of the *minimally preprocessed* data (Glasser et al., 2013) released within the HCP. Wilkins et al. (2014) explicitly state that their original data were not corrected for certain artifacts, and thus, generated data are affected correspondingly. Second, *Diffantom* implements the hindered and restricted compartments model (Assaf and Basser, 2005), which is a more complete model than the multi-tensor diffusion model.

A possible competitor to *Diffantom* is the phantom generated for the Tractography Challenge in ISMRM 2015. Similarly to *Diffantom*, the organizers used an HCP subject as source of structural information. While this phantom is designed for the bundle-wise evaluation of tractography (with the scores defined in the *Tractometer* (Côté et al., 2013), such as geometrical coverage, valid connections, invalid connections, missed connections, etc.), *Diffantom* is intended for the connectome-wise evaluation of results, yielding a tractography with a large number of bundles. Therefore, *Diffantom* and *FiberFox* are complementary as the hypotheses that can be investigated are different. Moreover, *Diffantom* does not require costly manual segmentation of bundles, highly demanding in terms of physiology expertise and operation time. The software workflow released with this data report (the *diffantomizer*) ensures the reproducibility of *Diffantom* and enables the generation of custom *diffantoms*. The *diffantomizer* is designed for, but not limited to, use HCP datasets as source of structural information.

### Conclusion

*Diffantom* is a whole-brain digital phantom generated from a dataset from the Human Connectome Project. *Diffantom* is presented here to be openly and freely distributed along with The *diffantomizer* workflow to generate new *diffantoms*. We encourage the neuroimage community to contribute with their own *diffantoms* and share them openly.

## DATA SHARING

The first *Diffantom* and its distorted version are available under the Creative Commons Zero licence (CC0) using the Dryad Digital Repository (reference here when published). The package is organized following the BIDS (Brain Imaging Data Structure, Gorgolewski et al. (2015)) standard. The associated software to “*diffantomize*” real dMRI datasets is available at https://github.com/oesteban/diffantom under an MIT license. *Phantomas* is available in https://github.com/ecaruyer/Phantomas under the revised-BSD license.

## AUTHOR CONTRIBUTIONS

All the authors contributed to this study. OE designed the data generation procedure, implemented the processing pipelines and generated the example dataset. EC implemented *Phantomas* (Caruyer et al., 2014), helped integrate the project with the simulation routines. OE, EC, AD thoroughly discussed and framed the aptness of the data in the community. AD, MBC, MJLC, and AS interpreted the resulting datasets. MBC, MJLC, and AS advised on all aspects of the study.

## ACKNOWLEDGMENTS

We thank Gert Wollny for his revision of this work. *Funding*: This study was supported by the Spanish Ministry of Science and Innovation (projects TEC-2013-48251-C2-2-R and INNPACTO XIORT), Comunidad de Madrid (TOPUS) and European Regional Development Funds, the Center for Biomedical Imaging (CIBM) of the Geneva and Lausanne Universities and the EPFL, as well as the Leenaards and Louis Jeantet Foundations.

## Footnotes

1 Available at http://www.tractometer.org/ismrm_2015_challenge/

2 Corresponding to the *5TT format* established with the latest version 3.0 of *MRTrix* (Tournier et al., 2012)

3 Since here we simulate healthy subjects, the last fraction map *T*_5_ is empty and can be omitted

## APPENDIX

Let 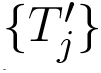 be the set of original fractions maps obtained with act `anat_prepare_fsl`, a tool in *MRTrix* that combines FAST (FMRIB’s Automated Segmentation Tool, Zhang et al. (2001)) and FIRST (FMRIB’s Integrated Registration and Segmentation Tool, Patenaude et al. (2011)) to generate the macrostructural 5TT map. FA denotes the fractional anisotropy (FA) map obtained from the original diffusion MRI (dMRI) data: the local fiber orientation maps {**V**_*i*_} with their estimated volume fractions 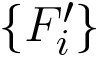 calculated with BEDPOSTX (Bayesian Estimation of Diffusion Parameters Obtained using Sampling Techniques modelling crossing –X– fibres, Jbabdi et al. (2012)). The final *T*_*j*_ maps of isotropic fractions are computed as follows:

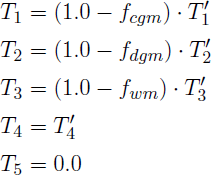

where *f* _{*cgm,dgm,wm*}_ are the fractions of restricted diffusion for each tissue. Sepehrband et al. (2015) found out that the fiber fraction ranges across the corpus callosum from the 70**±**8% in its body to an upper bound of 80**±**11% in the splenium. Therefore, we choose *f*_*wm*_ = 80% as default fraction of restricted diffusion in the white matter (WM). To our knowledge, restricted diffusion fractions have been studied only for WM. Therefore, we set *f*_*cgm*_ = 25% and *f*_*dgm*_ = 50% as they yield plausible FA and anisotropic diffusion coefficient (ADC) maps, assessed visually. The final {*F_i_*} maps are computed as follows:

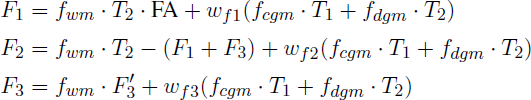

where *w*_{*f* 1,*f* 2,*f* 3}_ are the contributions of the gray matter (GM) compartments to each fiber population. By default: *w*_*f*1_ = 48%, *w*_f2_ = 37%, *w*_*f*3_ = 15%. Finally, the resulting maps are normalized to fulfill ∑*_j_ T_j_* + ∑*_i_ F_i_* = 1.0.

## REFERENCES

Alexander, A. L., Hasan, K. M., Lazar, M., Tsuruda, J. S., and Parker, D. L. (2001). Analysis of partial volume effects in diffusion-tensor MRI. Magn Reson Med 45, 770–780. doi:10.1002/mrm.1105

Assaf, Y. and Basser, P. J. (2005). Composite hindered and restricted model of diffusion (CHARMED) MR imaging of the human brain. NeuroImage 27, 48–58. doi:10.1016/j.neuroimage.2005.03.042

Basser, P. J., Pajevic, S., Pierpaoli, C., Duda, J., and Aldroubi, A. (2000). In vivo fiber tractography using DT-MRI data. Magn Reson Med 44, 625–632. doi:10.1002/1522-2594(200010)44:4 extless625::AID-MRM17 extgreater3.0.CO;2-O

Campbell, J. S. W., Siddiqi, K., Rymar, V. V., Sadikot, A. F., and Pike, G. B. (2005). Flow-based fiber tracking with diffusion tensor and q-ball data: Validation and comparison to principal diffusion direction techniques. NeuroImage 27,725–736. doi:10.1016/j.neuroimage.2005.05.014

Caruyer, E., Daducci, A., Descoteaux, M., Houde, J.-C., Thiran, J.-P., and Verma, R. (2014). Phantomas: a flexible software library to simulate diffusion MR phantoms. In 23th ISMRM (Milano, Italy), vol. 23

Caruyer, E., Lenglet, C., Sapiro, G., and Deriche, R. (2013). Design of multishell sampling schemes with uniform coverage in diffusion MRI. Magn Reson Med 69, 1534–1540. doi:10.1002/mrm.24736

Chua, T. C., Wen, W., Slavin, M. J., and Sachdev, P. S. (2008). Diffusion tensor imaging in mild cognitive impairment and Alzheimer’s disease: a review. Curr Opin Neurol 21, 83–92. doi:10.1097/WCO.0b013e3282f4594b

Close, T. G., Tournier, J.-D., Calamante, F., Johnston, L. A., Mareels, I., and Connelly, A. (2009). A software tool to generate simulated white matter structures for the assessment of fibre-tracking algorithms. NeuroImage 47, 1288–1300. doi:10.1016/j.neuroimage.2009.03.077

Côté, M.-A., Girard, G., Boré, A., Garyfallidis, E., Houde, J.-C., and Descoteaux, M. (2013). Tractometer: Towards Validation of Tractography Pipelines. Med Image Anal 17, 844–857. doi:10.1016/j.media.2013.03.009

Esteban, O., Daducci, A., Caruyer, E., O’Brien, K., Ledesma-Carbayo, M. J., Bach-Cuadra, M., et al. (2014). Simulation-based evaluation of susceptibility distortion correction methods in diffusion MRI for connectivity analysis. In 11th ISBI (Beijing, China), 738–741. doi:10.1109/ISBI.2014.6867976

Fieremans, E., De Deene, Y., Delputte, S., Özdemir, M. S., D’Asseler, Y., Vlassenbroeck, J., et al. (2008). Simulation and experimental verification of the diffusion in an anisotropic fiber phantom. J Magn Reson 190, 189–199. doi:10.1016/j.jmr.2007.10.014,

Fillard, P., Descoteaux, M., Goh, A., Gouttard, S., Jeurissen, B., Malcolm, J., et al. (2011). Quantitative evaluation of 10 tractography algorithms on a realistic diffusion MR phantom. NeuroImage 56, 220–234. doi:10.1016/j.neuroimage.2011.01.032

Garyfallidis, E., Brett, M., Amirbekian, B., Rokem, A., Van Der Walt, S., Descoteaux, M., et al. (2014). Dipy, a library for the analysis of diffusion MRI data. Front Neuroinform 8, 8. doi:10.3389/fninf.2014.00008

Glasser, M. F., Sotiropoulos, S. N., Wilson, J. A., Coalson, T. S., Fischl, B., Andersson, J. L., et al. (2013). The minimal preprocessing pipelines for the Human Connectome Project. NeuroImage 80, 105–124. doi:10.1016/j.neuroimage.2013.04.127

Golby, A. J., Kindlmann, G., Norton, I., Yarmarkovich, A., Pieper, S., and Kikinis, R. (2011). Interactive Diffusion Tensor Tractography Visualization for Neurosurgical Planning. Neurosurgery 68, 496–505. doi:10.1227/NEU.0b013e3182061ebb

Gorgolewski, K., Burns, C. D., Madison, C., Clark, D., Halchenko, Y. O., Waskom, M. L., et al. (2011). Nipype: a flexible, lightweight and extensible neuroimaging data processing framework in Python. Front Neuroinform 5, 13. doi:10.3389/fninf.2011.00013

Gorgolewski, K. J., Poline, J.-B., Keator, D. B., Nichols, B. N., Auer, T., Craddock, R. C., et al. (2015). Brain Imaging Data Structure - a new standard for describing and organizing human neuroimaging data. In INCF Neuroinformatics 2015 (Cairns, Australia). doi:10.3389/conf.fnins.2015.91.00056

Griffa, A., Baumann, P. S., Thiran, J.-P., and Hagmann, P. (2013). Structural connectomics in brain diseases. NeuroImage 80, 515–526. doi:10.1016/j.neuroimage.2013.04.056

Gössl, C., Fahrmeir, L., Pütz, B., Auer, L. M., and Auer, D. P. (2002). Fiber Tracking from DTI Using Linear State Space Models: Detectability of the Pyramidal Tract. NeuroImage 16, 378–388. doi:10.1006/nimg.2002.1055

Jbabdi, S., Sotiropoulos, S. N., Savio, A. M., Grafña, M., and Behrens, T. E. J. (2012). Model-based analysis of multishell diffusion MR data for tractography: How to get over fitting problems. Magn Reson Med 68, 1846–1855. doi:10.1002/mrm.24204

Jenkinson, M., Beckmann, C. F., Behrens, T. E., Woolrich, M. W., and Smith, S. M. (2012). FSL. NeuroImage 62, 782–790. doi:10.1016/j.neuroimage.2011.09.015

Kaus, M. R., Nabavi, A., Mamisch, C. T., Wells, W. H., Jolesz, F. A., Kikinis, R., et al. (2000). Simulation of Corticospinal Tract Displacement in Patients with Brain Tumors. In 3rd MICCAI (Pittsburgh, PA, US), LNCS 1935, 9–18. doi:10.1007/978-3-540-40899-4_2

Leemans, A., Sijbers, J., Verhoye, M., Van der Linden, A., and Van Dyck, D. (2005). Mathematical framework for simulating diffusion tensor MR neural fiber bundles. Magn Reson Med 53, 944–953. doi:10.1002/mrm.20418

Lin, C.-P., Tseng, W.-Y. I., Cheng, H.-C., and Chen, J.-H. (2001). Validation of Diffusion Tensor Magnetic Resonance Axonal Fiber Imaging with Registered Manganese-Enhanced Optic Tracts. NeuroImage 14, 1035–1047. doi:10.1006/nimg.2001.0882

Messé, A., Rudrauf, D., Giron, A., and Marrelec, G. (2015). Predicting functional connectivity from structural connectivity via computational models using MRI: An extensive comparison study. NeuroImage 111, 65–74. doi:10.1016/j.neuroimage.2015.02.001

Neher, P. F., Laun, F. B., Stieltjes, B., and Maier-Hein, K. H. (2014). Fiberfox: An Extensible System for Generating Realistic White Matter Software Phantoms. In MICCAI (Nagoya, Japan), vol. 22 of Mathematics and Visualization, 105–113. doi:10.1007/978-3-319-02475-2_10

Patenaude, B., Smith, S. M., Kennedy, D. N., and Jenkinson, M. (2011). A Bayesian model of shape and appearance for subcortical brain segmentation. NeuroImage 56, 907–922. doi:10.1016/j.neuroimage.2011.02.046

Perrin, M., Poupon, C., Rieul, B., Leroux, P., Constantinesco, A., Mangin, J.-F., et al. (2005). Validation of q-ball imaging with a diffusion fibre-crossing phantom on a clinical scanner. Philos T Roy Soc B 360, 881–891. doi:10.1098/rstb.2005.1650

Poupon, C., Rieul, B., Kezele, I., Perrin, M., Poupon, F., and Mangin, J.-F. (2008). New diffusion phantoms dedicated to the study and validation of high-angular-resolution diffusion imaging (HARDI) models. Magn Reson Med 60, 1276–1283. doi:10.1002/mrm.21789

Sepehrband, F., Clark, K. A., Ullmann, J. F., Kurniawan, N. D., Leanage, G., Reutens, D. C., et al. (2015). Brain tissue compartment density estimated using diffusion-weighted MRI yields tissue parameters consistent with histology. Hum. Brain Mapp. 36, 3687–3702. doi:10.1002/hbm.22872

Sporns, O., Tononi, G., and Kötter, R. (2005). The human connectome: A structural description of the human brain. PLoS Comput. Biol. 1, e42. doi:10.1371/journal.pcbi.0010042

Toda, K., Baba, H., Ono, T., and Ono, K. (2014). The utility of diffusion tensor imaging tractography for post-operative evaluation of a patient with hemispherotomy performed for intractable epilepsy. Brain Dev. 36, 641–644. doi:10.1016/j.braindev.2013.08.001

Tournier, J.-D., Calamante, F., and Connelly, A. (2012). MRtrix: Diffusion tractography in crossing fiber regions. Int J Imag Syst Tech 22, 53–66. doi:10.1002/ima.22005

Tournier, J.-D., Calamante, F., King, M., Gadian, D., and Connelly, A. (2002). Limitations and requirements of diffusion tensor fiber tracking: An assessment using simulations. Magn Reson Med 47, 701–708. doi:10.1002/mrm.10116

Tournier, J. D., Yeh, C.-H., Calamante, F., Cho, K.-H., Connelly, A., and Lin, C.-P. (2008). Resolving crossing fibres using constrained spherical deconvolution: Validation using diffusion-weighted imaging phantom data. NeuroImage 42, 617–625. doi:10.1016/j.neuroimage.2008.05.002

Tuch, D., Reese, T., Wiegell, M., Makris, N., Belliveau, J., and Wedeen, V. (2002). High angular resolution diffusion imaging reveals intravoxel white matter fiber heterogeneity. Magn Reson Med 48, 577–582. doi:10.1002/mrm.10268

Van Essen, D., Ugurbil, K., Auerbach, E., Barch, D., Behrens, T., Bucholz, R., et al. (2012). The Human Connectome Project: A data acquisition perspective. NeuroImage 62, 2222–2231. doi:10.1016/j.neuroimage.2012.02.018

Wassermann, D., Makris, N., Rathi, Y., Shenton, M., Kikinis, R., Kubicki, M., et al. (2013). On describing human white matter anatomy: the white matter query language. In 16th MICCAI (Nagoya, Japan), LNCS 8149, 647–654. doi:10.1007/978-3-642-40811-3_81

Wilkins, B., Lee, N., Gajawelli, N., Law, M., and Leporé, N. (2014). Fiber estimation and tractography in diffusion MRI: Development of simulated brain images and comparison of multi-fiber analysis methods at clinical b-values. NeuroImage doi:10.1016/j.neuroimage.2014.12.060

Zhang, Y., Brady, M., and Smith, S. (2001). Segmentation of brain MR images through a hidden Markov random field model and the expectation-maximization algorithm. IEEE Trans Med Imag 20, 45–57. doi:10.1109/42.906424

